# Induction of Ventral Spinal V0 Interneurons from Mouse Embryonic Stem Cells

**DOI:** 10.1101/2020.08.06.237115

**Authors:** Jennifer Pardieck, Manwal Harb, Shelly Sakiyama-Elbert

## Abstract

The ventral spinal population of V0 interneurons (INs) contribute to the coordinated movements directed by spinal central pattern generators (CPGs), including respiratory circuits and left-right alternation. One challenge in studying V0 INs has been the limited number of cells that can be isolated from primary sources for basic research or therapeutic use. However, derivation from a pluripotent source, such as has been done recently for other IN populations could reduce this issue. However, there is currently no protocol to specifically derive V0 interneurons from embryonic stem cells or induced pluripotent stem cells. To generate an induction protocol, mouse embryonic stem cells (mESCs) were grown in suspension culture and then exposed to retinoic acid (RA) and collected at different time points to measure mRNA expression of the V0 progenitor transcription factor marker, *Dbx1*, and post-mitotic transcription factor marker, *Evx1*. The cultures were also exposed to the sonic hedgehog signaling pathway agonist purmorphamine (purm) and the Notch signaling pathway inhibitor N-{N-(3,5-difluorophenacetyl-L-alanyl)}-(S)-phenylglycine-t-butyl-ester (DAPT) to determine if either of these pathways contribute to V0 IN induction, specifically the ventral (V0_V_) subpopulation. From the various parameters tested, the final protocol that generated the greatest percentage of cells expressing V0_V_ IN markers was an 8 day protocol using 4 days of suspension culture to form embryoid bodies followed by addition of 1 μM RA from days 4 to 8, 100 nM purm from days 4 to 6, and 5 μM DAPT from days 6 to 8. This protocol will allow investigators to obtain V0 IN cultures for use in *in vitro* studies, such as those examining CPG microcircuits, electrophysiological characterization, or even for transplantation studies in injury or disease models.

## Introduction

Spinal interneurons (INs) generate a complex relay between the body and the brain, as dorsal IN types confer sensory information while ventral IN types have roles in motor output. The ventral IN circuits formed through various interconnections with each other, motor neurons (MNs), and the dorsal IN types allow for the rhythmic, oscillatory movements – walking, breathing, swimming, etc. – that are generated from a circuit centralized in the spinal cord, the central pattern generator (CPG). These movements occur as neural circuits of the CPG excite or inhibit the appropriate muscle groups to create coordinated motions. Ventral INs thus must project axons in many directions along the rostro-caudal axis and relative to the midline, with commissurally-projecting INs involved in left-right coordination to allow for alternation or synchronous motion. V0 INs include a large proportion of cells having commissural axonal projections and are known to contribute to left-right alternation [1,2]. V0 IN progenitors (p0s) arise near the central-most ventral neural tube near the central canal and express the transcription factor Dbx1 [3] (Figure 1). These p0 progenitors mature into two major subclasses, ventral V0_V_ and dorsal V0_D_ INs, with the excitatory V0_V_ INs distinguished by transient expression of the transcription factor Evx1, while the inhibitory V0_D_ INs as yet have no specific, direct marker for their identification [1,4,5]. V0_V_ INs are further diversified into the Pitx2^+^ subclasses V0_G_ (glutamatergic) and V0_C_ (cholinergic), which are an uncommon, ipsilaterally-projecting V0 IN population that forms monosynaptic connections with MNs [6]. Genetic ablation studies in mice showed that the two major subclasses are recruited in a frequency-dependent manner during locomotion; the inhibitory V0_D_ INs are more active at low speeds and the excitatory V0_V_ interneurons at higher frequencies [2]. However, a recent study in larval zebrafish showed that, in contrast to mice, V0_D_ INs are important at higher frequencies [7]. Much remains to fully understand and characterize V0 IN subtypes and their role in locomotor circuits.

**Figure 1).**
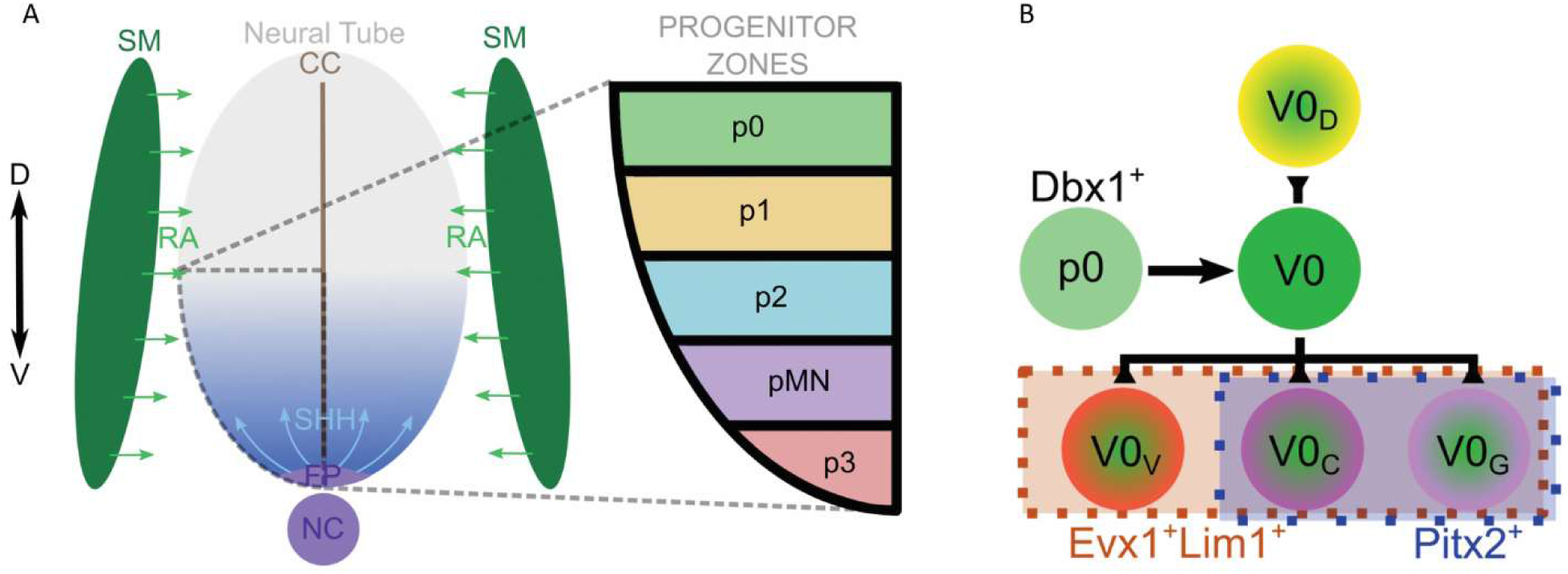
Schematic of RA and Shh gradient specifying IN progenitor domains and current distinctive markers of V0 IN subpopulations. (**A**) Release of retinoic acid (RA) from the adjacent somitic mesoderm (SM) specifies the spinal neural identity of the developing neural tube. Sonic hedgehog (Shh) released from the notochord (NC) and ventral (V) floor plate (FP) forms a concentration gradient; this gradient determines the different ventral interneuron progenitor zones for p3, pMN, p2, p1, and p0 which respectively become the V3 INs, MNs, V2 INs, V1 INs, and V0 INs; the dorsal (D) domains are not shown. (**B**) The V0 progenitors, p0, are identified by Dbx1 transcription factor expression and give rise to the post-mitotic V0 IN population. V0 INs have been defined by subpopulations of dorsal V0_D_, which currently have no distinct markers, and the ventral V0_V_ identified by expression of transcription factors Evx1 and Lim1. V0_V_ INs are further divided into the Pitx2-expressing glutamatergic V0_G_ and cholinergic V0_C_ populations.

To better study V0 IN populations and their contributions to CPGs, a means of obtaining a large number of V0 INs would be beneficial. Deriving INs from a pluripotent cell type, such as induced pluripotent stem cells (iPSCs) or embryonic stem cells (ESCs), is a means of isolating a potentially large number of INs from an expandable source. Recently, groups have used human iPSCs and mouse ESCs (mESCs) to generate different ventral IN populations including V1, V2a, and V3, as well as motor neurons (MNs) [8–13]. The work presented in this article shows the V0_V_ IN population can be differentiated from mESCs using similar methods as were previously successful in deriving ventral INs and MNs.

Approaches deriving INs and MNs from pluripotent sources recapitulate some aspects of normal spinal cord development. During this complex process, the ventral neural tube is exposed to retinoic acid (RA) from the adjacent somitic mesoderm and a gradient of sonic hedgehog (Shh) from a ventral source – beginning from the notochord and later including the floorplate (see Figure 1A). RA caudalizes the tissue, allowing for spinal identity versus midbrain or cortical identities, whereas the Shh gradient establishes the boundaries of the different IN progenitor domains. The progenitor and post-mitotic IN domains can be identified by particular transcription factor profiles, with progenitor V0 INs (p0s) expressing the homeobox protein Dbx1 [3], and a post-mitotic, ventral subpopulation – V0_V_ INs – expressing the homeobox protein Evx1 [1,4,5]. Lim1 is another transcription factor expressed in several ventral IN populations and distinguishes V0_V_ INs from a dorsal IN population, dI1, which transiently expresses Evx1 but not Lim1 [3]. Therefore, in this study, Dbx1 and Evx1 with Lim1 are used as markers of p0s and V0_V_ INs, respectively, in induced cultures derived from mESCs. A protocol to induce mESCs into INs should ideally produce a considerable proportion of INs among the heterogenous resultant mixture, yet practically, it should entail as simple and quick a procedure as possible.

To induce mESCs to form MNs and V2a INs, cells are cultured in suspension for 2 days in the absence of any morphogens (2^-^) and allowed to form embryoid bodies (EBs) to simulate embryogenesis. EBs are exposed to RA and a Shh agonist (purmorphamine [purm] for V2a and smoothened agonist [SAG] for MNs) are added to the EBs for another 4 days (4^+^). V2a IN induction also includes exposure to a Notch signaling inhibitor during the last 2 days to preferentially specify the V2a over V2b IN subtype. Of the ventral progenitor subtypes, p0s arise furthest from the floor plate, and, as it was previously shown that V0 INs can arise in the absence of Shh [3], it is plausible that V0 INs can be generated in the absence of Shh signaling factors; the p0 marker, transcription factor Dbx1, is also a class I transcription factor that is inhibited by Shh [14]. Therefore, when first developing the V0 IN induction protocol, conditions with exposure to different concentrations of RA without addition of Shh agonists (purm or SAG) were tested. After p0 and V0_V_ IN markers were detected in cultures only exposed to RA, activation of Shh signaling and inhibition of Notch signaling were also examined and were found to increase the proportion of cells expressing V0_V_ IN markers.

## Materials and Methods

### ESC maintenance

RW4 mouse embryonic stem cells (ATCC, SCRC-1018) were maintained in T-25 flasks coated with 0.1% gelatin (MilliporeSigma, G1393; in water) in complete medium containing 1000 U/mL leukemia inhibitory factor (LIF; MilliporeSigma, ESG1106) and 100 μM β-mercaptoethanol (BME; Thermo Fisher Scientific, 21985023) at 37°C in 5% CO_2_. mESCs were passaged by dissociating colonies with 0.25% trypsin ethylenediaminetetraacetic acid (trypsin-EDTA; Thermo Fisher Scientific, 25200072) for five minutes followed by quenching and trituration with excess complete medium. Single cells were plated in a new flask containing complete medium +LIF +BME at a 1:5 ratio for two days or until ∼80% confluent.

### Media formulations

Complete medium: Dulbecco’s Modified Eagle Medium (DMEM; Thermo Fisher Scientific, 11965; +L-Glutamine, high glucose) containing 10% newborn calf serum, 10% fetal bovine serum, 1x nucleoside solution (10 μM thymidine, and 30 μM of adenosine, cytosine, guanosine, and uridine).

DFK5 medium: DMEM/F12 (Thermo Fisher Scientific, 11320; +L-Glutamine, +Sodium Pyruvate, –HEPES, high glucose) containing 5% knockout serum replacement (Thermo Fisher Scientific, 10828), 1x Insulin, Transferrin, Selenium Solution (Thermo Fisher Scientific, 41400; 1.72 μM Insulin, 68.8 nM Transferrin, 38.7 nM Sodium Selenite), 0.5x non-essential amino acid solution (50 μM of each amino acid), 0.5x nucleoside solution (5 μM thymidine, 15 μM of adenosine, cytosine, guanosine, and uridine), and 55 μM BME.

Neuronal medium: DFK5 medium and Neurobasal medium (Thermo Fisher Scientific, 21103), 1:1 (v/v) ratio.

### V0_V_ interneuron induction and culture

Please note that the following protocol is the final protocol determined through the work presented in this publication. Please refer to the results section for descriptions of variations in testing protocol parameters.

RW4 mESCs were dissociated with 0.25% trypsin-EDTA, quenched with complete medium, and counted. 3 × 10^6^ single cells were pelleted at 300x*g*, the medium was aspirated, and the cells were suspended in 10 mL DFK5 medium in a 10 cm tissue culture-treated dish coated with 0.1% agar (in water) to allow for embryoid body (EB) formation. After 2 days (2^-^), EBs in DFK5 medium were collected into a 15 mL conical tube and allowed to settle for 10 minutes. The medium was aspirated, and 10 mL of fresh DFK5 medium was used to resuspend the settled EBs and return them to the agar-coated dish. After another 2 days (4^-^), ∼30 μL of EBs per cm^2^ were settled (e.g. for a 10 cm dish, settle 2.5 mL of EBs) in a 15 mL conical tube for 10 minutes. Old medium was aspirated and 10 mL of fresh DFK5 +1 μM all-*trans* retinoic acid (RA; MilliporeSigma, R2625: resuspended as a 20 mM stock in dimethyl sulfoxide [DMSO; MilliporeSigma, D2650]) +100 nM purmorphamine (purm; MilliporeSigma, 540223: resuspended as a 50 mM stock in DMSO) was used to resuspend settled EBs and plate them on a non-tissue culture-treated 10 cm dish coated with 0.1% gelatin. After 2 days (4^-^/2^+^), medium was aspirated and replaced with 10 mL of fresh DFK5 +1 μM RA +5 μM N-{N-(3,5-difluorophenacetyl-L-alanyl)}-(S)-phenylglycine-t-butyl-ester (DAPT; MilliporeSigma, D5942: resuspended as a 10 mM stock in DMSO). Induction was complete after another two days (4^-^/4^+^).

For cultures grown longer than 8 days, multi-well plates were coated with 0.01% poly-L-ornithine (MilliporeSigma, P3655; in 10 mM borate buffer, pH 8.3), rinsed three times with HEPES-buffered saline solution (HBSS, pH 7.2), and coated with 10 μg/mL laminin (natural mouse; Thermo Fisher Scientific, 23017015; in HBSS). Induced cultures were dissociated with 0.25% trypsin-EDTA, quenched with complete medium, counted, and plated on laminin-coated wells at a range of densities, depending on the end-point (to account for proliferation), to achieve ∼1 × 10^5^ cells/cm^2^ in neuronal medium supplemented with 1x B27 supplement, 1x GlutaMAX, and 5 ng/mL for each of brain-derived neurotrophic factor (BDNF, recombinant human, PeproTech, 450-02: resuspended as a 10 μg/mL stock in 0.1% bovine serum albumin [BSA, MilliporeSigma, A2058] in phosphate buffered saline [PBS]), glial cell line-derived neurotrophic factor (GDNF, recombinant human, PeproTech, 450-10: resuspended as a 10 μg/mL stock in 0.1% BSA in PBS), and neurotrophin-3 (NT-3, recombinant human, PeproTech, 450-03: resuspended as a 10 μg/mL stock in 0.1% BSA in PBS).

### Isolation of RNA, reverse transcription, and qPCR

To collect cultured cells for qPCR analysis, medium was aspirated and cells were detached by addition of 0.25% trypsin-EDTA followed by quenching and dissociation in complete medium. Cells were pelleted at 300x*g*. All medium was aspirated, and pellets were resuspended in RLT buffer as provided from the RNeasy Mini Kit (Qiagen, 74106). Pellets were either frozen at −80°C or immediately used with the RNeasy kit to isolate RNA per manufacturer instructions. 250 ng or 500 ng of RNA was reverse transcribed using the High-Capacity cDNA Reverse Transcription Kit (Thermo Fisher Scientific, 4368813) per manufacturer instructions.

For qPCR, a solution of ultrapure water, TaqMan Fast Advanced Master Mix (Thermo Fisher Scientific, 4444963), a TaqMan probe against mouse β-actin as a reference gene (Thermo Fisher Scientific, Mm02619580_g1, using VIC-MGB_PL dye), and the TaqMan probe against the target gene using FAM-MGB dye (*Dbx1*: Mm02344179_m1; *Evx1*: Mm00433154_m1) was prepared and loaded into MicroAmp Fast Optical 96-Well Reaction Plate (Thermo Fisher Scientific, 4346906) before loading each sample in triplicate. Plates were sealed, spun briefly to remove bubbles, and loaded into the QuantStudio 3 instrument for measurement. The fold change in mRNA expression levels were calculated using the comparative C_T_ method (2^^-ΔΔCT^ values) with β-actin as the reference gene relative to uninduced cultures as the reference sample.

### Immunocytochemistry and image analysis

Cultures were plated on 48-well, laminin-coated plates for immunocytochemistry (ICC) analysis. Day 8 time point cultures were plated for 2-4 hours before fixation, day 10 cultures were plated for 2 days and day 12 cultures for 4 days before fixation. Wells were rinsed once with PBS after aspirating the culture medium, then cells were fixed in 4% paraformaldehyde (PFA; MilliporeSigma, P6148) in 0.1M phosphate buffer for 20 minutes at room temperature. Cells were then exposed to 2% normal goat serum (NGS; MilliporeSigma, G9023) with 0.1% Triton X-100 (MilliporeSigma, X100; in PBS) to permeabilize and block for 30 minutes at room temperature. Primary antibodies (see Table 1) were diluted in 2% NGS with 0.1% Triton X-100 and cells were stained overnight at 4°C. Primary antibody solutions were removed and cells were washed 3 times for 10 minutes per wash with PBS. Secondary antibodies (see Table 2) were diluted in 2% NGS with 0.1% Triton X-100 and then filtered with a 0.22 μm PVDF syringe filter (MilliporeSigma, SLGV033RS). After adding secondary antibodies, plates were wrapped in foil and the cells were incubated for 1 hour at room temperature. Secondary antibody solutions were removed and cells were washed 3 times for 10 minutes per wash with PBS. Cells were then stained in 1:1000 Hoechst 33258 (Thermo Fisher Scientific, H3569; in PBS) for 10 minutes, rinsed once with PBS, then imaged with a DFC9000 GT camera (Leica) mounted on a DMi8 inverted widefield microscope (Leica) using a SOLA Light Engine light source (Lumencor). Images were analyzed using a CellProfiler [15,16] pipeline to determine the percentage of cells co-stained for βIII-tubulin, Evx1, and Lim1. At least 2 images each were taken from at least 2 wells for each condition at each time point with N = 3-6.

**Table 1:** Primary antibodies used for ICC and flow cytometry.

| Target | Final concentration | Source, product number |
| --- | --- | --- |
| Evx1 | 0.5 µg/mL | Developmental Studies Hybridoma Bank, 99.1-3A2<sup>a</sup> |
| Evx1 | 1 µg/mL | Biorbyt, orb183448<sup>b</sup> |
| Lim1 | 1<sup>a</sup>, 5<sup>b</sup> µg/mL | Developmental Studies Hybridoma Bank, 4F2 |
| βIII tubulin | 1 µg/mL | Biolegend, 845501<sup>a</sup> |
| βIII tubulin | 0.5<sup>a</sup>, 1<sup>b</sup> µg/mL | Biolegend, 802001 |

<sup>a</sup>Used for ICC
<sup>b</sup>Used for flow cytometry

**Table 2:**
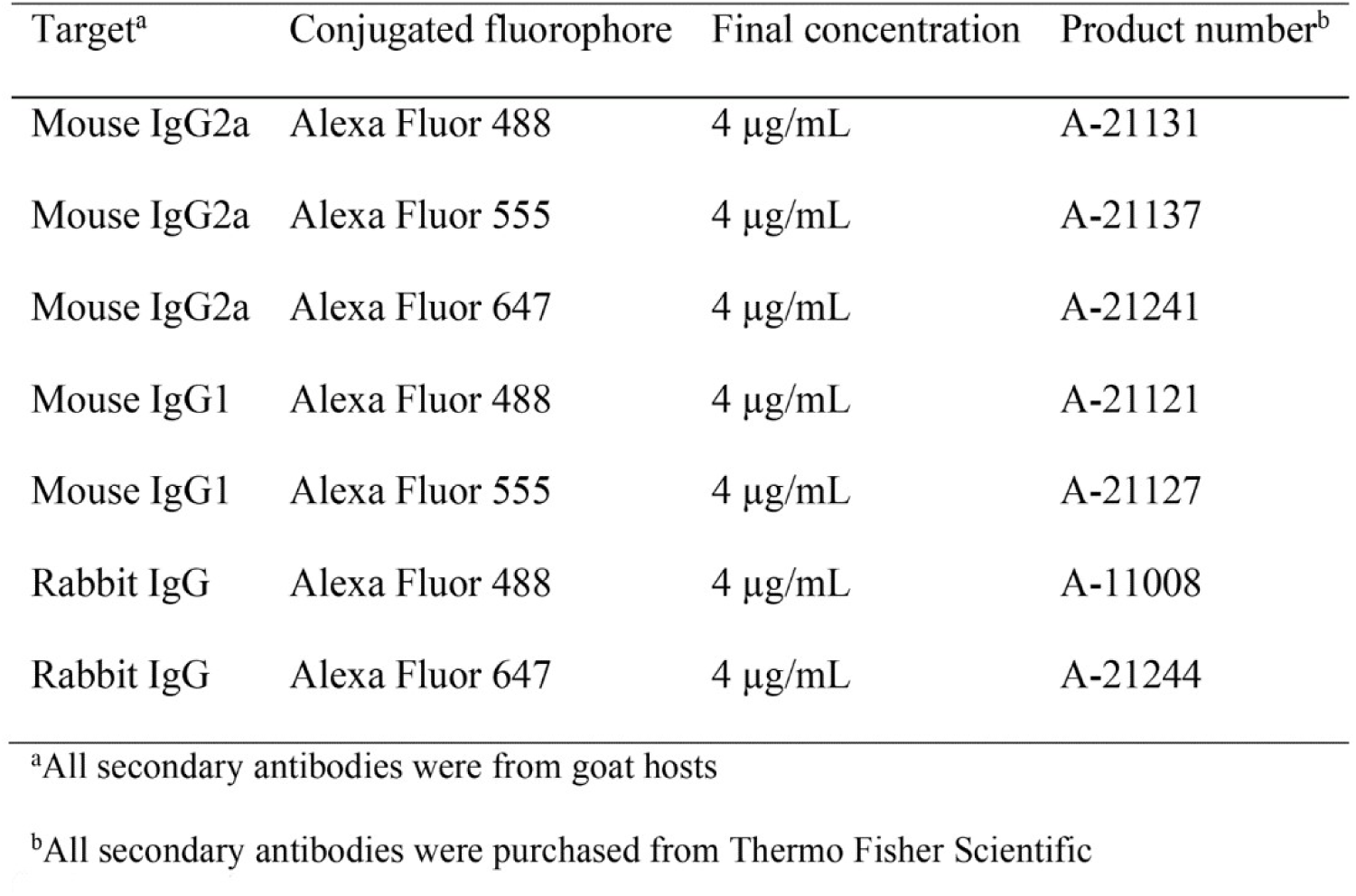
Secondary antibodies used for ICC and flow cytometry.

### Flow Cytometry

Cultures were plated on 6-well, laminin-coated plates for flow cytometry analysis. Before collection, day 8 time point cultures were plated for 2-4 hours, day 10 cultures for 2 days and day 12 cultures for 4 days. For collection, medium was aspirated and cells were detached by addition of 0.25% trypsin-EDTA followed by quenching and dissociation in complete medium. Cells were spun at 400x*g* for 5 minutes, medium aspirated, and pellets were resuspended in 4% PFA in 0.1M phosphate buffer to fix for 10 minutes at room temperature. For staining, samples were divided and pelleted at 400x*g*. For staining with Evx1 and Lim1 antibodies, samples were resuspended in 5% normal NGS with 0.1% Triton X-100 (% v/v;) in PBS to permeabilize and block for 20 minutes at room temperature. For staining with βIII tubulin antibody, samples were resuspended in 5% NGS with 0.1% saponin (% w/v; MilliporeSigma, S4521) in PBS to permeabilize and block for 20 minutes at room temperature. Primary antibodies (see Table 1) were diluted in 2% NGS in PBS and cells were stained for 1 hour at room temperature. Primary antibody solutions were removed and cells were washed 3 times for 10 minutes per wash with PBS. Secondary antibodies (see Table 2) were diluted in 2% NGS with in PBS and then filtered with a 0.22 μm PVDF syringe filter. After adding secondary antibodies, samples were protected from light while incubating for 1 hour at room temperature. Secondary antibody solutions were removed and cells were washed 3 times for 10 minutes per wash with PBS. Cells were resuspended in PBS for measurement on an Attune NxT Flow Cytometer (Thermo Fisher Scientific).

### Statistics

GraphPad Prism version 7 and Microsoft Excel were used for statistical analyses. Outliers were identified and excluded. Values are reported as means and error bars are standard error of the mean (S.E.M.). One-way analysis of variance (ANOVA) using Scheffe’s multiple comparison method with 95% confidence was used to determine significance, which is indicated in figures as follows, unless otherwise stated: * for p < 0.05, ** for p < 0.01, *** for p < 0.001, and **** for p < 0.0001.

## Results

### Generating V0 progenitor and IN marker-expressing cells from mESCs

With the goal of achieving a protocol specific to deriving V0 INs from mESCs, different induction conditions were tested by modifying procedures known to produce V2a INs and MNs [9,12]. By first culturing embryoid bodies (EBs) through growing mESCs in suspension for either 2 or 4 days (known as 2^-^ or 4^-^, see Figure 2A), embryogenesis is stimulated as well as the beginning of lineage specification, including neurogenesis. The EBs are then exposed to RA, which is involved in inducing neuralization and in caudalization towards a spinal fate. Several concentrations of RA were tested at amounts known to induce spinal neurons [12,17,18]. To determine whether these conditions effectively generated cells expressing V0 progenitor and IN markers, cultures were collected for qPCR analysis. As seen in Figure 2B, 2 days of EB formation followed by 4 days of exposure to RA resulted in a significant decrease in *Dbx1* (progenitor) mRNA expression at all concentrations tested except 4 μM RA and little change in *Evx1* (V0_V_ INs) mRNA expression over uninduced cultures grown under the same conditions without RA exposure. However, allowing 4 days of EB formation produced a significant increase in the *Dbx1* mRNA expression versus uninduced cultures after exposure to 1 or 2 μM RA for 2 days (4^-^/2^+^, Figure 2C). Another 2 days of RA exposure (4^-^/4^+^) generated a significant increase in *Evx1* mRNA expression over uninduced cultures at 1 μM RA (Figure 2D). Based on these data, a 4^-^/4^+^ protocol using 1 μM RA was used in further protocol development.

**Figure 2).**
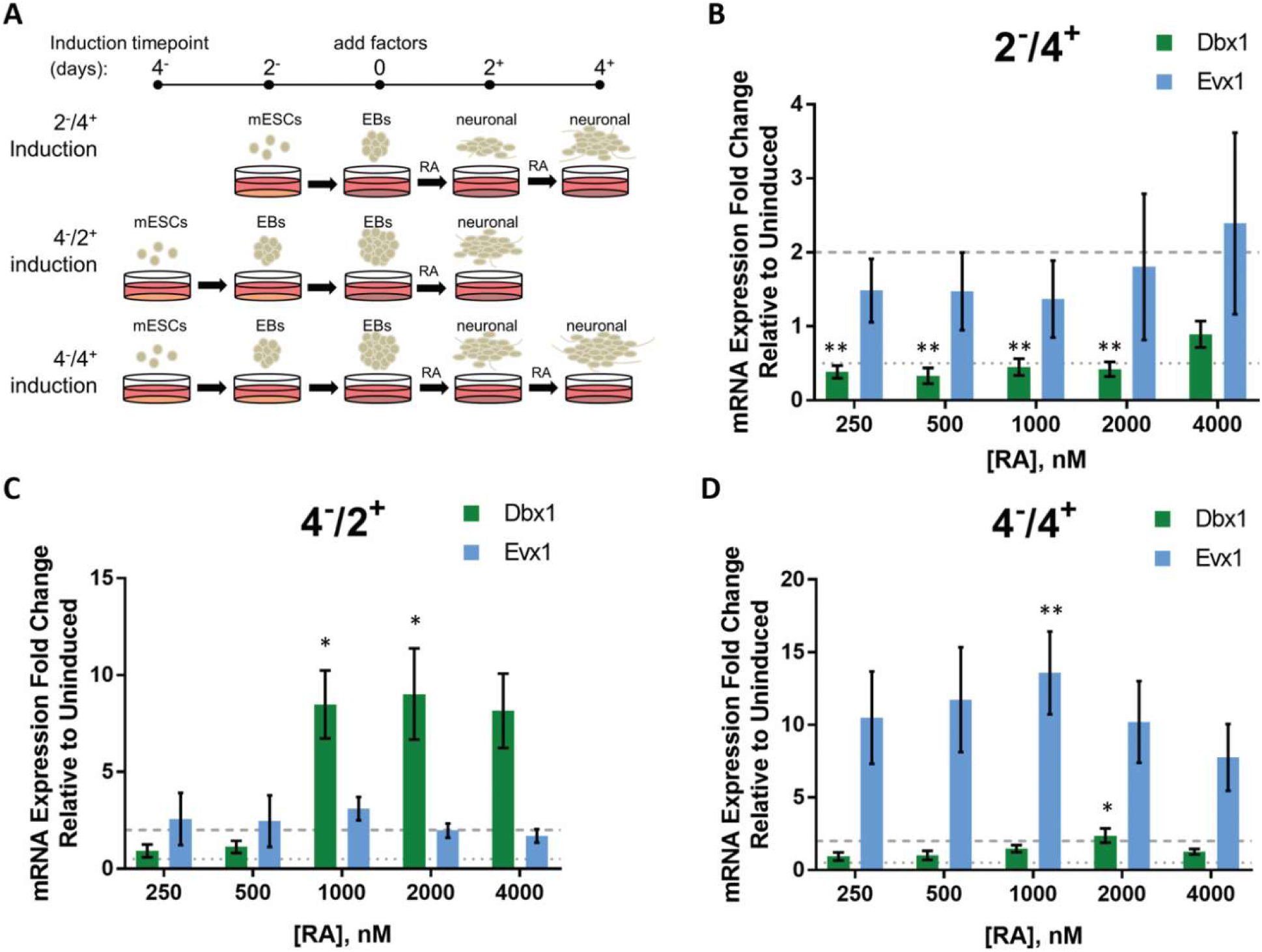
Determining the time course for induction of V0 progenitor and IN markers in vitro. **(A)** Schematic of the time course for induction of mESCs toward V0 IN fate. mESCs were suspended in agar-coated plates to allow formation of EBs for either 2 (2^-^) or 4 (4^-^) days. EBs were then plated on gelatin and exposed to RA for either 2 (2^+^) or 4 (4^+^) days before collection for qPCR analysis. **(B - D)** mRNA expression of progenitor marker *Dbx1* (green bars) and post-mitotic V0_V_ subtype marker *Evx1* (blue bars) in cells collected after inductions using various concentrations of RA for **(B)** 2^-^/4^+^, N = 4-11; **(C)** 4^-^/2^+^, N = 4-14; or **(D)** 4^-^/4^+^, N = 5-18. Values are given as fold change relative to uninduced cultures grown under the same conditions but without exposure to RA; --- line delimits upregulation, … line delimits downregulation. Error bars are S.E.M. Significance is reported for the same gene at the same time point: * denotes p < 0.05 vs uninduced, ** denotes p < 0.01vs uninduced.

### Improving yield of V0_V_ IN subtype

Although V0 progenitor and IN markers were induced when cultures were exposed only to RA, we wanted to determine the effect of Shh signaling, as it is known to be involved in ventral progenitor IN domain specification. Also, within the different IN populations, there exist many subtypes where the specification of some subtypes (i.e. V2a versus V2b) depends in part on Notch signaling. Therefore, to see if Shh and Notch signaling had an effect on the specification of V0_V_ INs, various conditions were tested with small molecule effectors for these pathways. The weak smoothened agonist purmorphamine (purm) was used to stimulate Shh signaling while the gamma-secretase inhibitor DAPT was used to impede Notch signaling. Initially, 4^-^/2^+^ cultures induced with 1 μM RA were examined to see the effects of different purm concentrations on V0 progenitor marker *Dbx1* mRNA expression; 10 nM and 100 nM purm did not significantly affect *Dbx1* mRNA expression, while 1 μM purm, a concentration used to induce V2a INs [9] from mESCs, significantly decreased *Dbx1* mRNA expression (Figure 3A). To examine Shh signaling in V0_V_ IN induction, 4^-^/4^+^ cultures induced with 1 μM RA were exposed to 100 nM purm for different intervals (see Figure 3B). Some of these cultures were also treated with 5 μM DAPT at the latter part of induction to determine whether Notch plays a role in V0_V_ IN induction as well. Figure 3C shows that exposure to 100 nM purm from days 4 to 6 or 5 to 8 with addition of DAPT from days 6 to 8 resulted in the greatest increase in *Evx1* mRNA expression (64.9 fold or 60.8 fold over uninduced cultures, respectively). These conditions also resulted in significantly greater *Evx1* mRNA expression over induction with RA and DAPT without purm, suggesting that affecting both Shh and Notch signaling is important for achieving a greater level of V0_V_ IN induction. Based on qPCR data of the conditions tested, the most efficient method of inducing *Evx1* mRNA expression from mESCs is using a 4^-^/4^+^ protocol with addition of 1 μM RA from days 4 to 8, 100 nM purm from days 4 to 6, and 5 μM DAPT from days 6 to 8.

**Figure 3).**
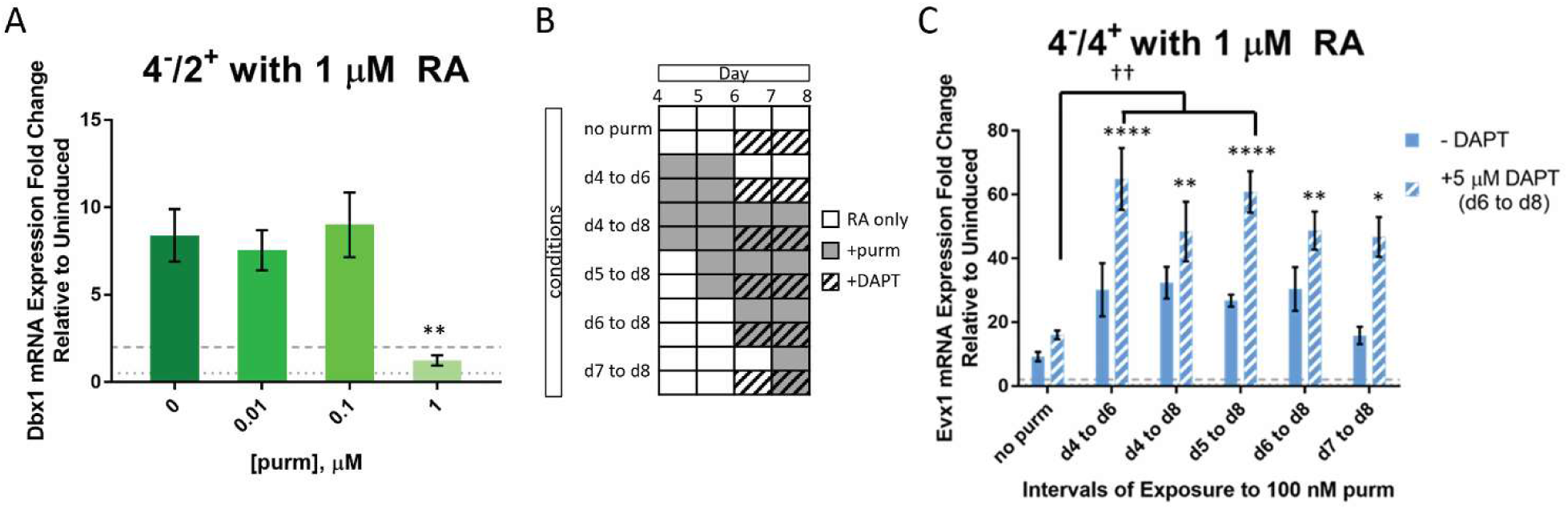
Inducing Shh signaling and inhibiting Notch signaling increases *Evx1* mRNA expression. mESCs were allowed to form EBs for 4 days. Cultures were then grown on gelatin and exposed to 1 μM RA from day 4 to day 8. **(A)** At day 4, inductions were tested with/without exposure to log-fold concentrations of purm up to 1 μM, which is known to induce V2a INs in vitro. Cultures were analyzed at 4^-^/2^+^ for *Dbx1* mRNA expression. N = 8-9. **(B)** Schematic of conditions tested to examine *Evx1* mRNA expression. At day 4, inductions were tested with/without exposure to 100 nM purm at various time intervals and with/without 5 μM DAPT from days 6 to 8. **(C)** *Evx1* mRNA expression evaluated at day 8. N = 4-11. For **(A)** and **(C)**, values are given as fold change relative to uninduced cultures grown under the same conditions but without exposure to RA; --- line delimits upregulation, … line delimits downregulation. Error bars are S.E.M. and significance is reported as follows: * denotes p < 0.05 vs only RA, ** denotes p < 0.01 vs only RA, and **** denotes p < 0.0001 vs only RA; †† denotes p < 0.01 for indicated samples vs induction with RA and DAPT.

### Quantifying proportion of cells expressing markers for V0_V_ INs at different time points

To ensure induction of cells expressing V0_V_ IN markers, we examined cultures by ICC with antibodies against Evx1, Lim1, and the pan neuronal marker βIII tubulin. Lim1 was used to distinguish the Lim1^+^ V0_V_ INs from a dorsal IN population, dI1, which also transiently expresses Evx1 during development but is Lim1^-^ [3]. Uninduced cultures, 4^-^/4^+^ cultures with only RA, and 4^-^/4^+^ cultures with RA, purm, and DAPT were stained after dissociation and plating on laminin-coated plates. Cell were plated at lower densities for day 10 and day 12 cultures to control for cell proliferation; however, final cell densities were still variable and often higher at the later time points. These cultures were fixed on days 8, 10, and 12: Figure 4A shows representative images of each condition and time point. Images were analyzed using a CellProfiler pipeline to determine the proportion of “triple positive” cells expressing βIII tubulin, Evx1, and Lim1 (Figure 4B). Triple positive cells were greatest in day 10 cultures grown with RA, purm, and DAPT, with approximately 44% of cells expressing V0_V_ IN markers. As an additional measurement of the proportion of cells expressing V0_V_ IN markers, we stained uninduced cultures and 4^-^/4^+^ cultures exposed to only RA or RA, purm, and DAPT on days 8, 10, and 12 and analyzed them by flow cytometry. Flow data show that addition of RA, purm, and DAPT significantly increases the percentage of cells expressing Evx1 and Lim1 at days 10 and 12 over uninduced cultures. At day 10, ∼57% of RA, purm, DAPT-induced cells express Evx1 and Lim while 60% of cells are positive for βIII tubulin – this proportion is comparable to the values seen by ICC analysis.

**Figure 4).**
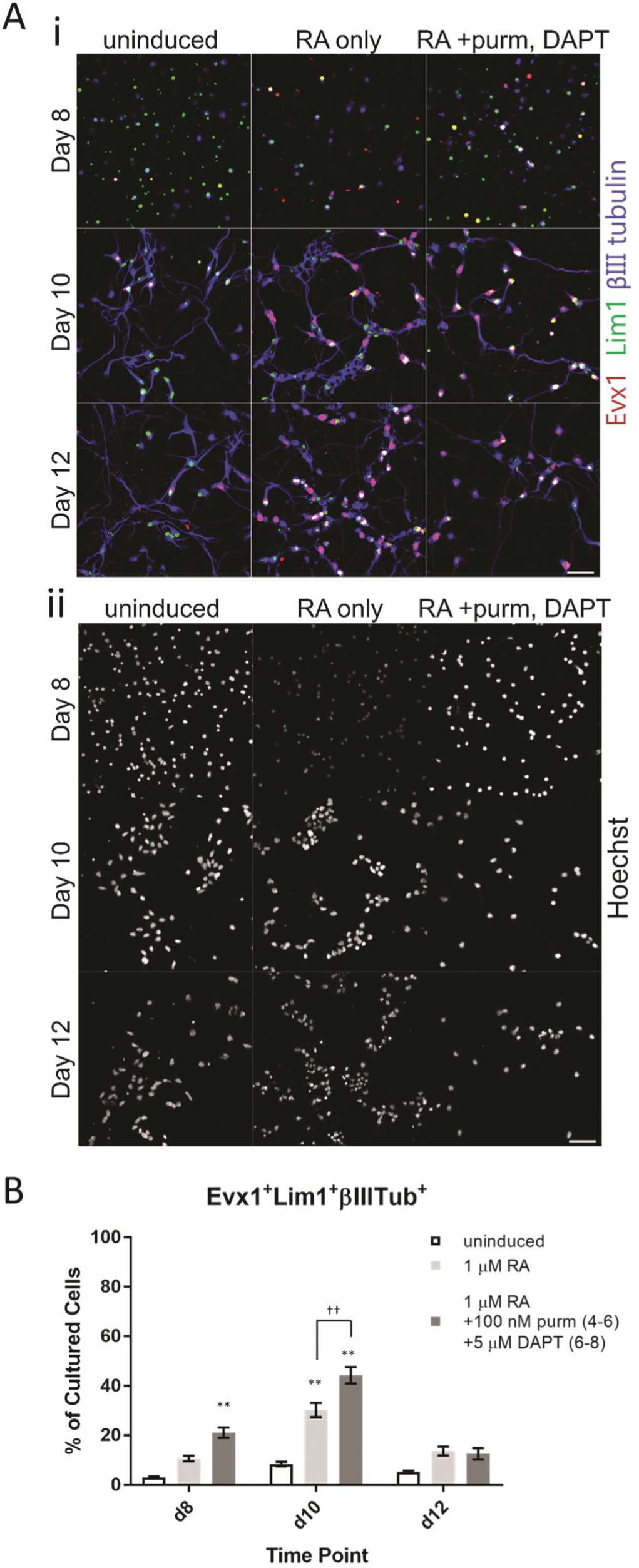
Immunocytochemistry shows day 10 cultures induced with RA, purm, and DAPT have the greatest proportion of cells expressing V0_V_ IN markers. mESCs were induced with 4^-^ /4^+^ protocol and fixed on day 8, day 10 or day 12; cultures were dissociated on day 8 and thereafter grown on laminin-coated plates in neuronal medium until fixation 2-4 hours, 2 days, or 4 days later for day 8, day 10, and day 12 time points, respectively. **(A)** Representative images of day 8, 10 and 12 cultures that were uninduced; exposed to 1 μM RA only; or exposed to 1 μM RA, 100 nM purm from days 4 to 6 and 5 μM DAPT from days 6 to 8. (**i**) Staining for Evx1 is shown in red, Lim1 in green, and βIII tubulin in blue. (**ii**) Hoescht was used to stain nuclei. Scale bars are 50 μm. **(B)** Percentages are given for the proportion of “triple positive” cells for each condition on days 8, 10 and 12. Error bars are S.E.M. and at least 2 images of at least 2 technical replicates were taken for each biological replicate: N = 3-6, n = 16-35. Significance is reported among samples of the same time point: ** denotes p < 0.01 vs uninduced; †† denotes p < 0.01 for RA, purm, and DAPT vs only RA.

**Figure 5).**
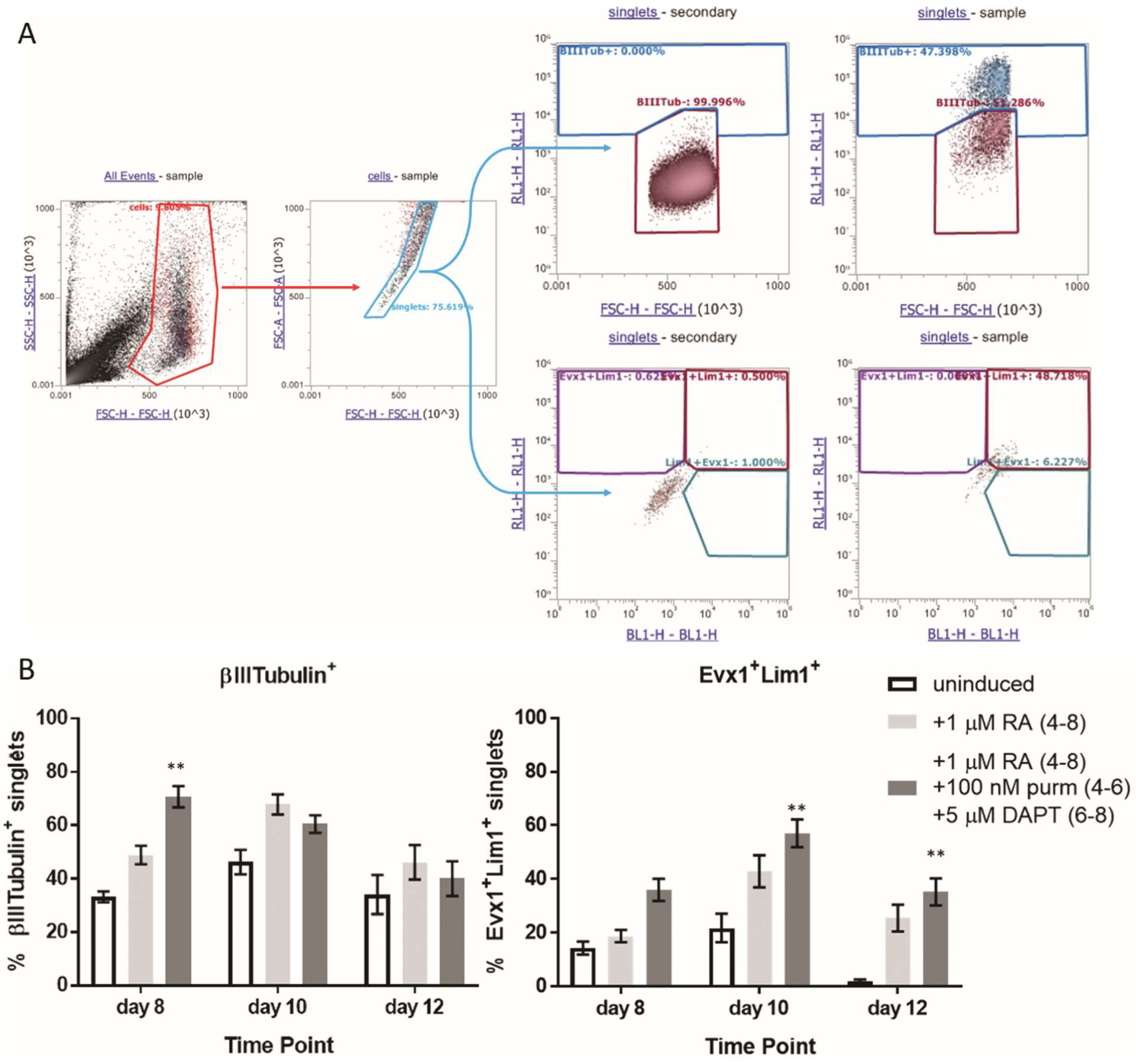
Flow cytometry shows day 10 cultures induced with RA, purm, and DAPT have the greatest percentage of Evx1^+^Lim1^+^ cells. mESCs were induced with 4^-^/4^+^ protocol, then dissociated and fixed on day 8, day 10 or day 12. Cultures were dissociated on day 8 and thereafter grown on laminin-coated plates in neuronal medium until fixation 2-4 hours, 2 days, or 4 days later for day 8, day 10, and day 12 time points, respectively. **(A)** Schematic of gating strategy. The “cells” gate identifies the IN cell population versus debris. Singlets are gated to exclude cell clumps. Secondary only-stained cells are used to gate singlets that do not stain for βIII tubulin or Evx1 and Lim1. **(B)** Percentages are given for the proportion of βIII tubulin^+^ cells or for Evx1^+^Lim1^+^ cells for each condition on days 8, 10 and 12. Error bars are S.E.M. For βIII tubulin staining, N = 8-9. For Evx1 and Lim1 staining, N = 11-13. Significance is reported among samples of the same time point: ** denotes p < 0.01 vs uninduced.

## Discussion

The objective of this study was to derive V0 INs from mESCs with a simple and short protocol. While other inductions deriving MNs or IN populations from mESCs have used 2 days for EB formation prior to exposure to morphogens, this study has shown that allowing 4 days of EB formation produces higher V0 progenitor and V0_V_ IN marker mRNA expression. Perhaps the additional 2 days provides a necessary increase in cell layers in the EBs to better simulate the conditions of V0 progenitor development *in vivo* (positional depth in the tissue in combination with morphogen concentration), as these cells are known to arise near the medial central canal.

Compared to established protocols to induce ventral neuronal populations, similar concentrations of RA led to significant increases in V0 marker mRNA expression, with 1 μM RA producing the greatest effect for inducing both the progenitor marker *Dbx1* and the V0_V_ IN marker *Evx1*. Although exposure to only RA was sufficient to induce V0_V_ IN marker *Evx1* mRNA expression, providing additional factors to stimulate Shh signaling and inhibit Notch signaling improved the proportion of cells expressing V0_V_ IN markers significantly (Figure 3C). For Shh, the less potent purm was chosen over SAG to achieve signaling levels closer to those known to induce V2a INs. It seems there is a threshold concentration for purm to affect V0 progenitor development, as the 10 and 100 nM concentrations had little effect on *Dbx1* mRNA expression over induction with only RA; exposure to 1 μM purm significantly reduced *Dbx1* mRNA expression (Figure 3A). This idea is supported by the fact that 1 μM purm is the concentration used to induce mESCs to form V2a INs^8^, a more ventral population exposed to a higher concentration of Shh during specification. Also, previous studies have shown and discussed that threshold effects contribute to the definition of the progenitor boundaries [14,19–22]. Based on this effect seen on *Dbx1* mRNA expression, further studies were completed using 100 nM purm, the highest concentration tested without effect on *Dbx1* mRNA expression. It is possible that induction of cells expressing V0_V_ IN markers would be enhanced if further testing were completed to find the best concentration of purm for induction. Different time courses of Shh stimulation through purm exposure showed that earlier addition at days 4 or 5 tended to yield higher *Evx1* mRNA expression. However, addition of purm alone did not significantly increase marker expression over induction with only RA; addition of the Notch inhibitor DAPT alone from days 6 to 8 also did not significantly increase *Evx1* mRNA expression over RA only cultures. When both are present during the induction, however, there is a synergistic effect producing significantly higher expression in all cultures exposed to both factors relative to inductions only exposed to RA. Here, a later addition of DAPT was tested, as subtype specification has been suggested to occur later in the developmental timeline [23], as occurs with V2a versus V2b INs [24,25]. Other time courses of DAPT exposure might alter *Evx1* mRNA expression levels, but a brief examination of addition of 5 μM DAPT from days 4 to 6 was also tested with no obvious changes over RA-only induction in the proportion of V0_V_ INs stained by ICC (data not shown).

The data presented in this study have shown that the derived cells include those that express V0_V_ IN markers; however, other cell types are also induced in these cultures, which might obscure any observations on the function of V0_V_ INs obtained from utilizing the induced cultures in further studies. Therefore, a means of purification of V0_V_ INs from the heterogeneous population is desirable. A high-purity culture of V0_V_ INs would provide a tool for investigators for use in studies including analyzing spinal microcircuits in a controlled manner, such as in microelectrode arrays or microdevices, or after transplantation in animal models to determine whether V0_V_ INs contribute to any functional recovery post-SCI. Recently, neural progenitor cells and high-purity V2a INs were transplanted in a cervical level contusion injury rat model, and rats receiving the V2a INs showed increased functional recovery over other groups [26]; this shows the utility of obtaining a large, high-purity population of INs and the promise of their therapeutic potential. This study has provided the first step for future investigations using isolated V0_V_ INs: a method of obtaining a high proportion of V0_V_ INs from a renewable source with a simple 8 day induction protocol.

With a protocol available to derive a large number of V0_V_ INs from a renewable source lies an opportunity for enabling future investigations. Researchers can study the interactions between particular types of INs in vitro by plating populations of interest in microdevices or multi-electrode arrays to observe the interactions between the different cell types, thus contributing to our knowledge of the roles of INs in the CPG. V0_V_ INs can also easily be cultured at a high enough number after only 8 days for use in animal transplantation studies to determine their role in recovery after loss of function due to disease or injury. Other spinal populations, MNs and V2a INs for example, have been derived from mESCs as well as from human pluripotent cells [13,27]; V0_V_ INs could potentially be derived from human pluripotent cells using a modified version of this protocol, bringing this research a step closer to translational therapeutics.

## Abbreviations

IN: interneuron
CPG: central pattern generators
mESCs: mouse embryonic stem cells
RA: retinoic acid
purm: purmorphamine
DAPT: N-{N-(3,5-difluorophenacetyl-L-alanyl)}-(S)- phenylglycine-t-butyl-ester
V0V: ventral V0
MN: motor neuron
p0: progenitor V0 interneuron
V0D: dorsal V0
V0G: glutamatergic V0
V0C: cholinergic V0
iPSC: induced pluripotent stem cell
Shh: sonic hedgehog
EBs: embryoid bodies
SAG: smoothened agonist
LIF: leukemia inhibitory factor
BME: β-mercaptoethanol
EDTA: ethylenediaminetetraacetic acid
DMEM: Dulbecco’s Modified Eagle Medium
HBSS: HEPES-buffered saline solution
BDNF: brain-derived neurotrophic factor
BSA: bovine serum albumin
PBS: phosphate buffered saline
GDNF: glial cell line-derived neurotrophic factor
NT-3: neurotrophin-3
ICC: immunocytochemistry
PFA: paraformaldehyde
NGS: normal goat serum
SM: somitic mesoderm
NC: notochord
V: ventral
D: dorsal

## Acknowledgements

The authors were funded by NIH R01NS090617 (SSE) and F31NS100432 (JP). The authors acknowledge Nisha Iyer, Hao Xu, Sarah Oswald, Mary Salazar, and Nick White for guidance, discussion, and technical assistance.

